# BioGeoFormer: A deep learning approach to classify unknown genes associated with critical biogeochemical cycles

**DOI:** 10.64898/2025.12.17.695047

**Authors:** Jacob H. Wynne, Nima Azbijari, Andrew R. Thurber, Maude M. David

## Abstract

Remote functional annotation continues to impede progress in microbial ecology, as alignment-based approaches still leave over one-third of microbial sequences functionally unresolved. In contrast, pre-trained natural-language–processing approaches have shown strong potential for inferring functions from diverse biological sequences, and here we introduce a protein language modeling approach allowing us to classify sequences into 37 defined key pathway categories involved in 4 major biogeochemical cycles (methane, sulfur, nitrogen and phosphorus cycles). To do so, we fine-tuned ESM2-8m using databases curated for biogeochemical cycling pathways. Our resultant **BioGeo**chemical cycling trans**Former** (BioGeoFormer or BGF) was high-performing on validation and test sets, producing embeddings that exhibit an ability to infer protein function at a metabolic pathway level. BGF was applied to a dataset of metagenome-assembled genomes (MAGs) constructed from methane-fueled, deep-sea, “cold seep” environments to demonstrate its utility in contrast to current informatics approaches. We employed multiple gene assignments to identify gene function within these MAGs. A total of 1.05M genes were assigned biogeochemical functions, with BGF alone suggesting putative ecosystem roles for 0.49M (46%) of these at a confidence of 85% or greater; these genes were classified as unknown by the other approaches. Across the pathways of interest, BGF identified 6 times as many genes, on average, as Hidden Markov models (HMMs) as well as alignment-based approaches across the various pathways. BGF provides a novel tool that is capable of informing process-based hypotheses in diverse systems, highlighting cryptic proteins most notably linked to methane, nitrogen, and phosphorus cycling while uncovering the mysteries within microbial dark matter.

**Author summary:** When investigating the function of microbes in the environment, scientists are often left with vast amounts of genes or proteins where no knowledge about their function is available. This represents a huge amount of information left to be discovered in many fields of biology. One recent approach that has shown significant potential in further understanding unknown proteins are protein language models, which are deep-learning methods leveraging large datasets to understand the ‘language’ of proteins. We aimed to apply protein language modeling to further understand the function of proteins as they relate to large-scale environmental transformation of nutrients and carbon. Specifically, we designed our approach to further understand the metabolism of microbes that affect methane, nitrogen, phosphorus, and sulfur, all elements that are highly impactful to the planet’s function and health. We used our new approach on a deep-sea microbiology dataset, and showed the method’s utility in further understanding the function of proteins and their impact on the environment. Overall, we found our method is an important new tool in the toolset of environmental scientists working to better understand the function of microbes and their proteins.

## 1. Introduction

Microbial communities are the driving force of many global biogeochemical cycles, including carbon [1], nitrogen [2], sulfur [3], and phosphorus [4]. Over the past two decades, advances in high-throughput sequencing have led to a global sampling of microbial community structure and helped advance the mechanistic understanding of microbiome driven earth-system processes. Widespread application of metagenomic techniques have resulted in large-scale discoveries of microbial genomes across most of the world’s biomes [5], but our ability to link genes, genomes, and microbial community structure to their underlying processes, is severely impeded by the significant proportion of gene sequences without a known function [6]. This gap in understanding of the microbial world is colloquially referred to as “microbial dark matter” (MDM). Sampling of MDM via metagenomics and environmental DNA (eDNA) is among the most rapidly expanding “big data” sets globally [7], however while we are producing massive amounts of sequence data much of it falls within the category of “unknown”. MDM represents an estimated 1 million microbial taxa and 1 billion genes that have no known identity or function [6].

Protein functional annotation represents a crucial and ever-evolving forefront of this challenge. To date the most widespread method to predict protein function is through sequence alignment or identity searches (e.g., BLAST, DIAMOND), where an input sequence is compared to a database of sequences with previously annotated functions [8]. Additionally, Hidden Markov models [9] are also a gold standard for protein sequence annotation, offering greater sensitivity than pairwise alignment methods by leveraging information from multiple sequence alignments (MSAs) [10].

Although pairwise and profile alignment approaches have contributed significantly to function prediction, they only allow reliable inference of function for high levels of sequence identity (greater than 60%) and there is significant bias towards function described in highly studied organisms [8]. This limitation is exacerbated by the fact that annotated proteins and organisms are those that lend themselves to culturing and experimentation, which precludes as much as 99% of the microbial taxa in marine environments [11]. As stochastic models, HMMs represent a hidden sequence of states that probabilistically emit observable symbols, capturing the statistical properties of sequence families [12]. In practice, a separate HMM is constructed for each functional group — typically defined by shared annotations—using an MSA of representative sequences. Consequently, generating *N* HMM profiles for *N* distinct protein functions requires computing *N* corresponding MSAs. This approach is computationally intensive and also depends heavily on the quality of the MSAs—something that becomes particularly challenging for proteins with unknown functions, which often lack well-annotated homologs or sufficient sequence diversity to support robust profile construction [13]. Yet, there exists a large pool of functionally analogous proteins that are remotely homologous to known orthologs or functionally convergent [13,14].

An emergent approach to address these limitations of alignment and HMM methods is application of deep learning approaches, especially language models. Deep learning and natural language processing (NLP) algorithms can identify and extract complex, latent features and patterns from data. Their use in protein function prediction have increased dramatically over the last decade [15] in part as a result of the large amount of data produced during the metagenomics revolution [16]. Deep learning language models using transformer architecture in particular have been shown to go beyond capturing sequence variation and also encode information about diverse biological properties [17–19]: these representations have been shown to support accurate prediction of protein structure and subsequently function [20,21].

In this study, we introduce a novel framework that combines two approaches: 1) using pre-trained transformer models and 2) detecting new orthologs using entire pathways as classification categories (instead of individual proteins). We first propose to leverage pre-trained transformer models trained on large-scale biological sequence datasets. As stated above, transformers exhibit the ability to learn generalizable features that capture other properties than sequence identity, such as structural, functional, and evolutionary patterns across diverse taxa [19]. Further, *pre-trained* transformer models, such as ESM [19], ProtBERT [22], and AlphaFold [21], are especially well-suited for annotating Microbial Dark Matter because they have been shown to encode sequence information in a way that supports effective transfer learning, enabling robust performance even with limited task-specific labeled data [19,21,22]. By utilizing these pre-trained protein representations, we can also substantially reduce the computational burden associated with training deep models from scratch and accelerate the development of predictive tools for protein function, structure, and interaction.

Second, we extended our analysis beyond individual protein annotations and focused on categorical grouping of proteins per pathway rather than individual protein annotation for protein discovery. Recent studies have successfully used this approach and yielded functional discoveries by leveraging protein language models in antibiotic resistance [23], prokaryotic viral protein function [24,25], and phosphorous cycling protein families [26]. Our database proposes to combine these approaches (pathway classification on pre-trained models) specifically tuned on key biogeochemical pathways that affect planetary health which are still lacking in the literature.

Understanding biogeochemical cycling and the microbial processes that shape them is critically important across scientific domains, including climate [27,28], agriculture [29], biotechnology [30], ecosystem function [31,32], and human health [33] among others. Developing computational techniques to better understand microbial mechanisms as they relate to biogeochemical cycling have critically lagged compared to domains such as antibiotic resistance [34,35] and gut-microbiome functions [36–38].

Our study proposes to address these gaps, and fine tunes the model ESM2-8m using 1.9 million proteins from 4 manually curated and published databases containing metabolic pathways linked to four biogeochemical cycles especially relevant to climate, as well as chemosynthetic and photosynthetic primary production: methane, sulfur, nitrogen, and phosphorus [20,39–41]. Proteins were subdivided into 37 classes defined by metabolic pathways. We then applied our BioGeochemical transFormer (BioGeoFormer or BGF) to a comprehensive catalog of metagenome-assembled genomes constructed from cold seeps around the globe [42], an environment characterized by diverse microenvironments, microbial communities, and biogeochemical cycles, all associated with substantial unexplored genetic novelty.

## 2. Results

### 2.1. Development, optimization and performance evaluation of BioGeoFormer across identity gradients

#### 2.1.1. A transformer-based classifier was developed and tested on environmental sequence datasets partitioned at multiple sequence-identity thresholds

BioGeoFormer (BGF) was optimized on 4 curated databases comprising 3.2 M protein sequences [20,39–41] (Fig 1), which were filtered to 1.9 M pathway-specific sequences. Briefly (see detailed descriptions of the procedures in the Methods section), the database was constructed by combining four previously published datasets, which were manually curated across 37 pathways and filtered to include only proteins belonging to one pathway with the exception of the central methanogenic pathway and anaerobic oxidation of methane (Table S1) (we refer to this database as BioGeoFormer-db (BGFdb)). Using CD-HIT, each class within BGFdb was clustered at distinct sequence similarities from 20% to 90% at an interval of 10%, and then split into a 60/20/20 scheme for training, validation, and test data respectively. We stringently filtered the test set using DIAMOND alignment post clustering, ensuring sequences were not aligned at a higher percent identity to the training dataset than their respective splits (Figs S1-S4). We used ESM-2, a protein language model pre-trained on the protein sequence database UniRef (8 million parameter transformer with 6 layers, named ESM2-8m), and added a classification layer of 37 units (corresponding to our curated cycles), allowing us to exploit the evolutionary representations that ESM-2 has learned from millions of sequences.The model was entirely re-trained for each data identity split. Results were benchmarked against DIAMOND (using individuals pairwise comparison to the training datasets) and Hidden Markov Models that were trained on all the sequences for each pathways (35-37 models for each identity split).

**Fig 1.**
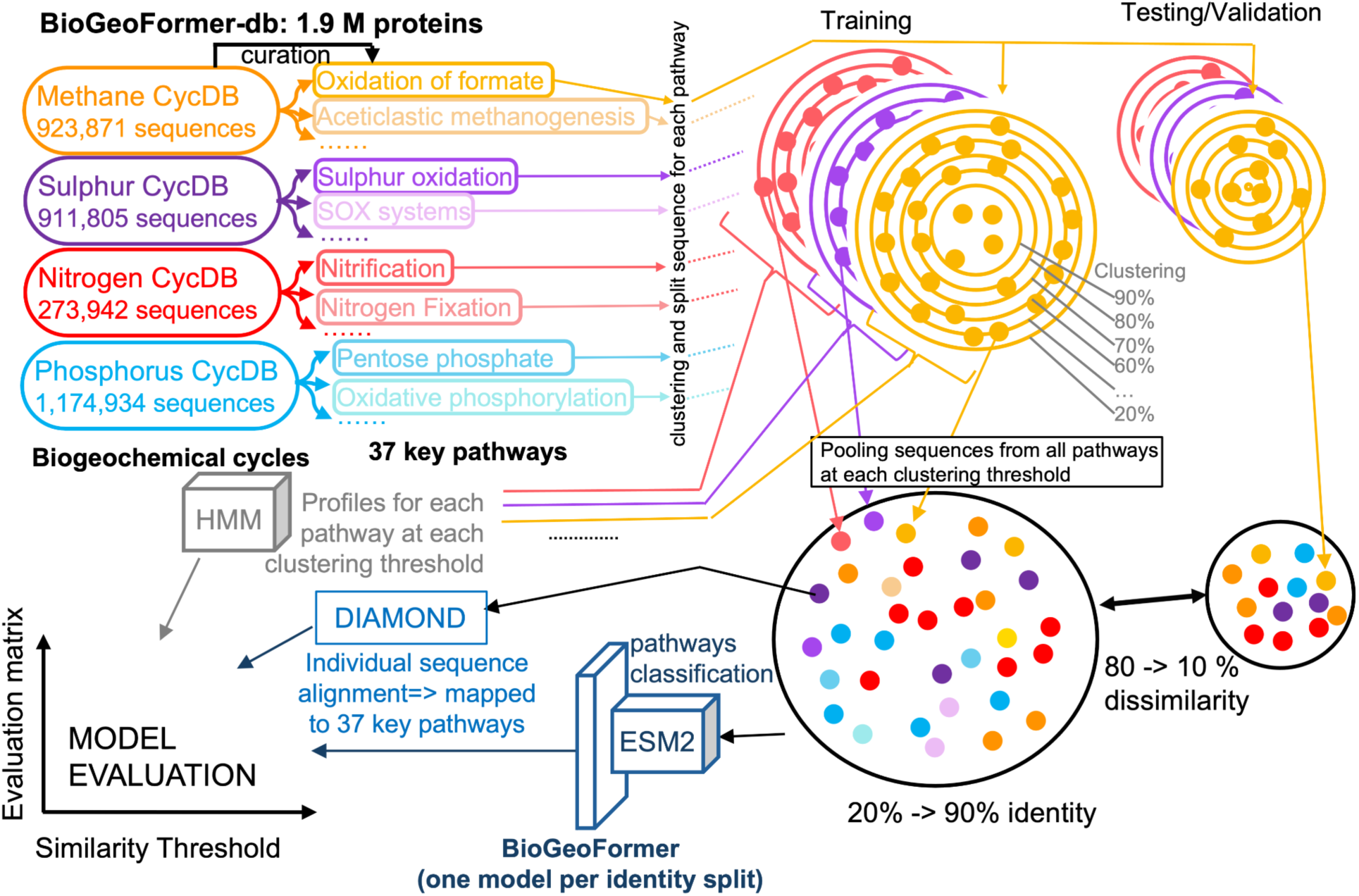
Pipeline for development and evaluation of BioGeoFormer’s (BGF) performance, and benchmark against gold standard methods (DIAMOND, and HMMs). Data was curated from MCycDB (orange), SCycDB (purple), NCycDB (red), PCycDB (blue) and processed according to methods discussed in this study. To evaluate BGF at identifying distant, yet functionally related proteins we clustered using CD-HIT from 90-20% identity, between training, validation, and test sets. Clustering was done per pathway to ensure for each cycle, providing an approach to testing how far the model can generalize outside of its training data.

#### 2.1.2. BGF returned high-performing scores across all identity splits and performance metrics for validation and showed a significant correlation between identity split and performance on the filtered-test set

We aimed to quantify the ability of the three approaches (BGF, HMMs and DIAMOND) to identify proteins on the testing set (Table 1). In this case, appended test sets (see: Methods: data curation) had proteins removed that were more similar than the identity score being considered; for example at 20% all proteins that were 20.1% similar or greater were removed from the test and validation sets prior to attempting to classify them. Results in table one demonstrate performance of each model split applied to their respective filtered-test sets (test sets were filtered to avoid data leakage inherent to CD-HIT; *see methods: dataset curation*). Performance ranged from 94-95% when considering all metrics for the 90% identity split, and 9-15% for the 20% identity split (maintaining a 80% identity distance between testing and training).

**Table 1.**
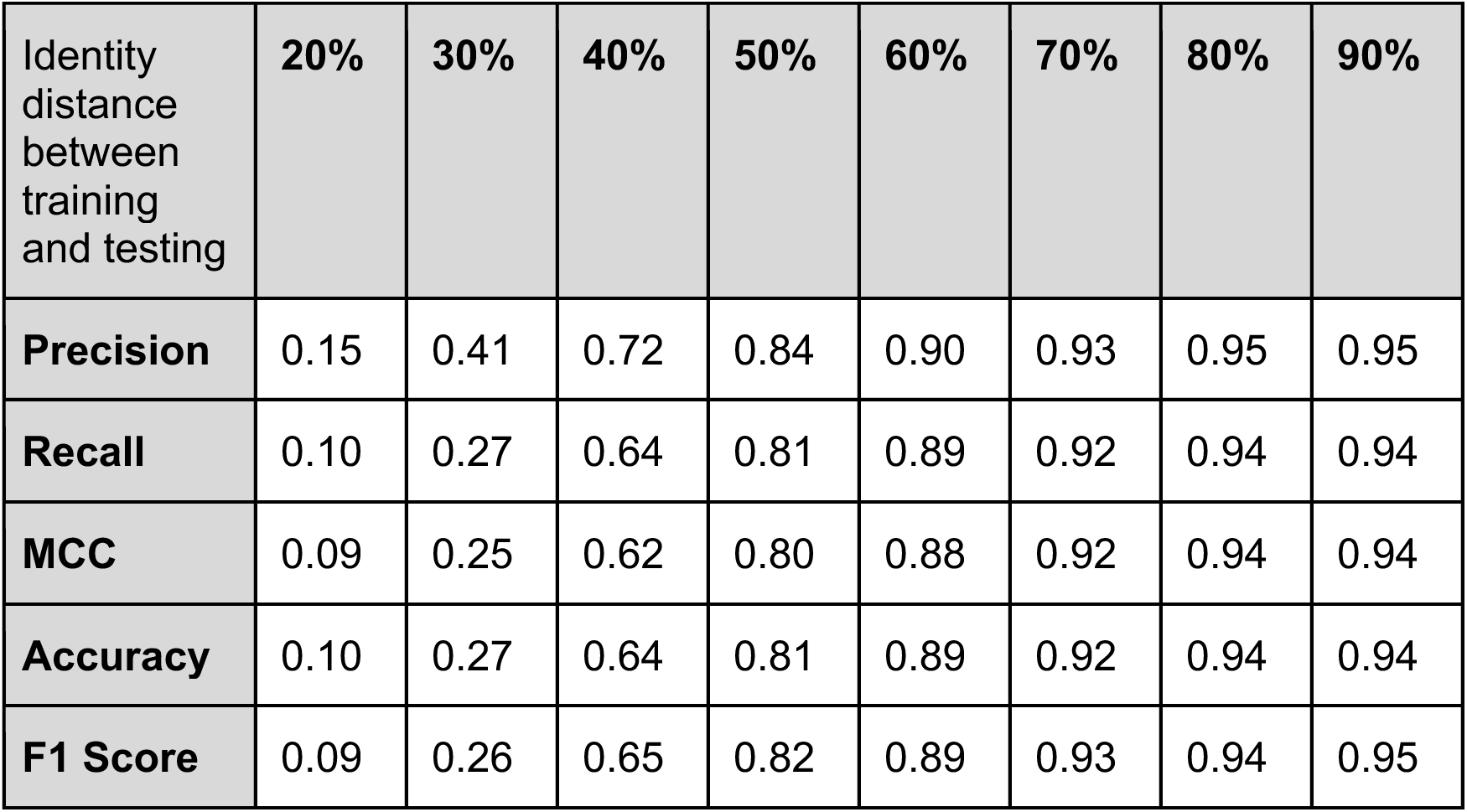
Performance metrics of BioGeoFormer (BGF) against filtered-test corresponding to each identity split.

A limitation of the accuracy metric is that it can misrepresent model performance when trained or tested on unbalanced class sizes associated with the various cycles, in contrast the Matthew’s Correlation (MCC) can better quantify model performance [43]. MCC supported the trends of the accuracy metric in this case, remaining within 5%, even though the cycles identified were unbalanced in their representation. BGF’s precision exceeded its recall, indicating that the model produced relatively few false positives, but it missed some true positives, especially significant at distant splits (e.g., precision was 41% while recall was 27% at the 30% split).

#### 2.1.3. BGF outperformed hidden markov models trained on the same datasets at all identity thresholds, with significant improvements in detecting true-positives, and outperformed DIAMOND at distant identity thresholds

BGF performed comparably or better than alignment based approaches and better than Hidden Markov Models (HMMs) across the range of identity splits. Unsurprisingly, when comparing alignment using DIAMOND between known dataset clusters and training and test sets there was a high level of success across all metrics (Fig 2), including outperforming BGF at high identity splits. HMMs, in contrast, performed poorer than BGF and DIAMOND, with an inflection point at 50% identity within the models, demonstrating a reduced ability of true positives at higher identity scores, a possible sign of overfitting. At the lowest identity split (20%), we found that BGF outperforms both alignment and HMMs, highlighting the potential for BGF to functionally identify proteins at remote homologies. The most illustrative point is that at 20% identity, BGF’s MCC was 0.09 and precision was 0.15 in contrast to 0.01 and 0.04 for HMMs, respectively and 0.01 and <0.001 for DIAMOND (Fig 2a,e). At this extreme case, this 9 fold increase in MCC and more than 3 fold increase in precision in contrast to the other methods suggests a potential for putative classification of very poorly constrained genes. As identity increases, BGF continues to consistently outperform HMMs across all metrics, however as training and test sets become more similar, HMMs increase in their ability to characterize the genes in the test and validation sets.

**Fig 2.**
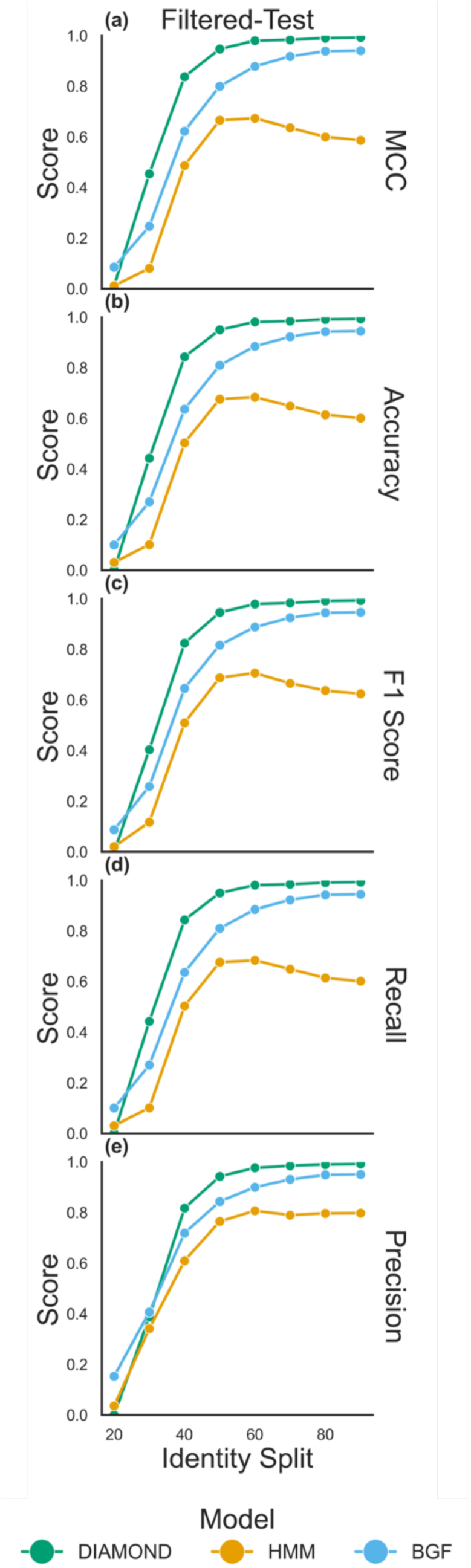
Performance metrics of BGF (blue), HMM models (yellow), and DIAMOND (green) trained at different sequence identity splits. Every point represents a unique model trained and evaluated on a filtered-test set split based on sequence identity splits (*see methods: dataset curation*). Metrics of performance include Matthews Correlation Coefficient (MCC) (**a**), accuracy (**b**), F1 score (**c**), recall (**d**), and precision (**e**) and are discussed at further length in the methods (*see methods: performance metrics*). Evaluation is carried out on the eight BGF models, DIAMOND alignments on each training split, and the 35-37 HMMs per identity split.

#### 2.1.4. BGF showed lower prediction error for pathway assignments at higher sequence identity levels, with nitrification as a weak point across both near and distant identity

BGF was able to correctly classify the majority of proteins across the 37 cycles considered, however its performance across cycles was uneven (Fig 3). Overall, between 20% and 90% the percent of miscategorized proteins decreased from 16.1% to 5.5%. However, proteins belonging to some specific cycles, like nitrification and sulphur oxidation were consistently misclassified. Even at the 90% split, Nitrification was miscategorized as dissimilatory sulphur reduction and oxidation (dsro) proteins, organic phosphate hydrolysis (org_phos_hyd), and methane oxidation.

**Fig 3.**
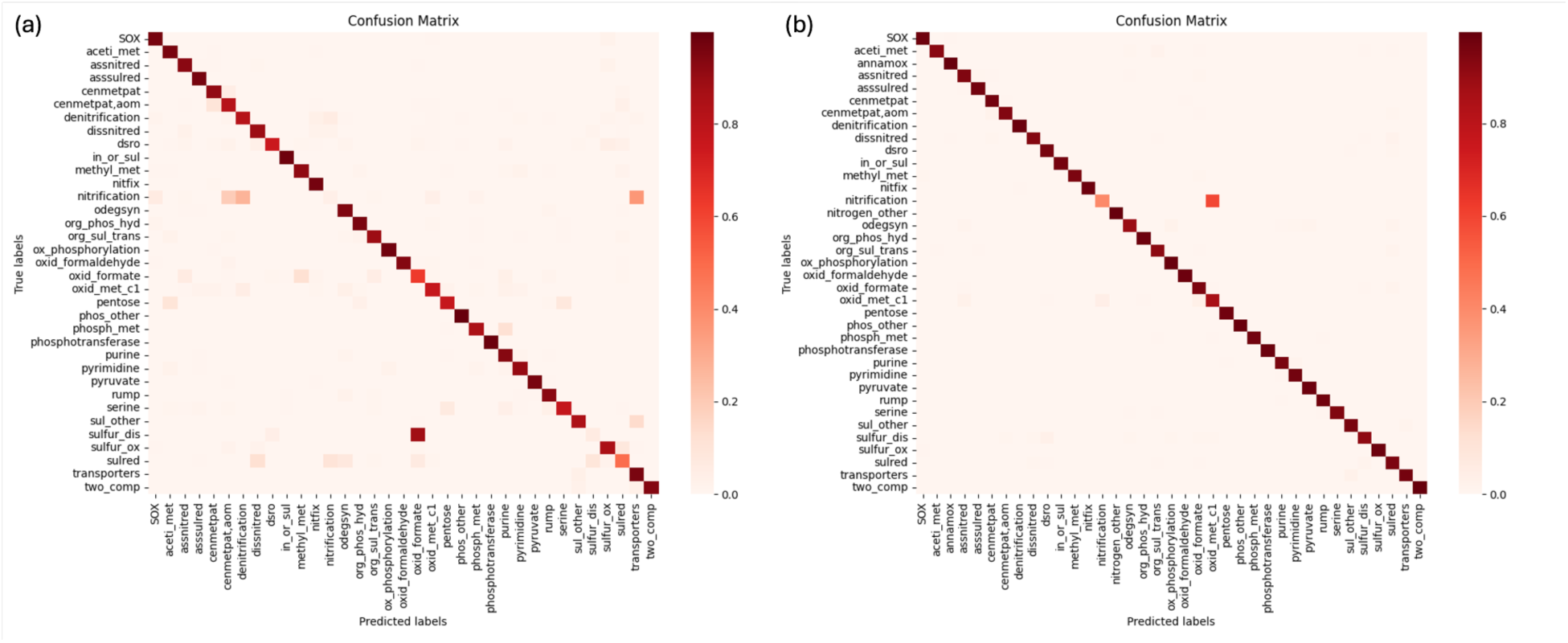
Confusion matrices for 20% (**a**) and 90% (**b**) sequence identity split model outputs on their respective test sets. Darker regions off of the diagonal represent misannotations (a.k.a. confusion) by each model comparing true labels (y-axis) to predicted labels (x-axis)

#### 2.1.5. T-SNE representation of embeddings produced by BGF

The t-SNE representation of the last hidden layer of BGF illustrates the model is learning a representation that captures the biogeochemical cycles when split at 20%. (Fig 4a), as well as, for comparison, 90% identity to the training set (Fig 4b). Overall, between both of these data sets, we observed clear separation for the majority of the proteins classified, regardless of the identity among data sets employed (Figs 4, S5-S10). While there are still some proteins with ambiguous function, as identified by color overlap (Fig 4), these results highlight the potential importance of BGF to classify protein data sets that are poorly represented in training sets.

**Fig 4.**
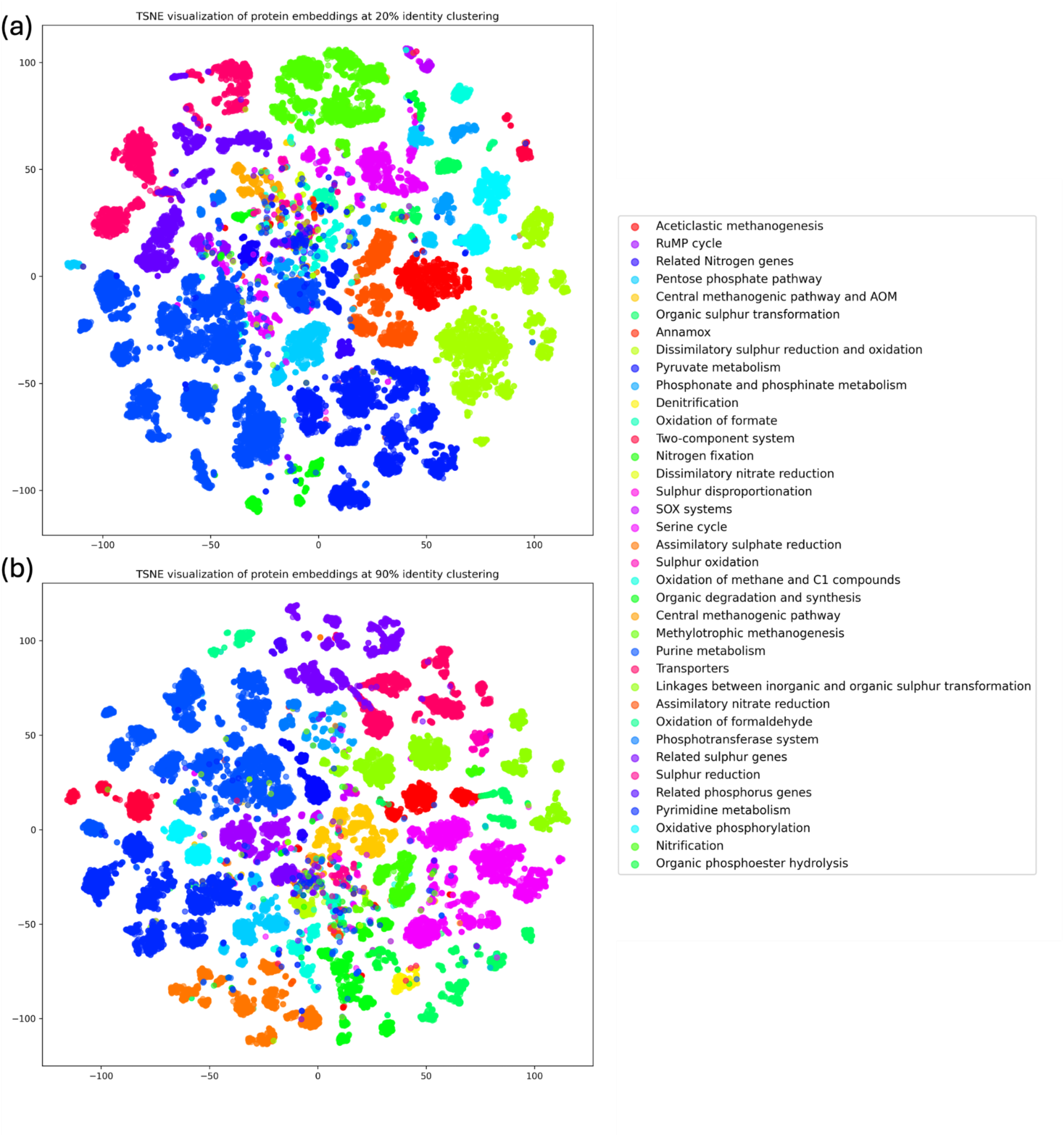
T-distributed Stochastic Neighbor Embedding (t-SNE) visualization of randomly sampled clustered embeddings by BGF at 20% (**a**) and 90% (**b**) sequence identity splits for test sets. t-SNE plots provide 2D representation of high dimensional data, in this case demonstrating BGF’s ability to distinguish and group proteins by their biogeochemical cycle. Coloration of points are proteins designated based on the biogeochemical pathways in the BGFdb database (*see methods: dataset curation*). For each plot, 30,000 sequences were randomly selected from their respective test sets, perplexity was set to 30, and the learning rate was set to ‘auto’ with 1000 iterations.

### 2.2. Calibrating confidence and application on a deep sea dataset

#### 2.2.1 Calibration by temperature scaling led to more representative confidence distributions, with varying results across identity splits

We used the validation set to calibrate BGF, allowing us to mitigate model overconfidence and increase the accuracy of BGFs predictions. To achieve this, temperature scaling was used to calibrate confidence values for the softmax output, leading to more representative distributions of accuracy. Using negative log-likelihood (NLL; Fig 5a), expected calibration error (ECE; Fig 5b), and reliability diagrams, we assessed the success of temperature scaling under different conditions (*see methods: transformer model calibration*). We found that prior to temperature scaling, models were highly overconfident as shown by high NLL, high ECE and reliability diagrams (Fig 5).

**Fig 5.**
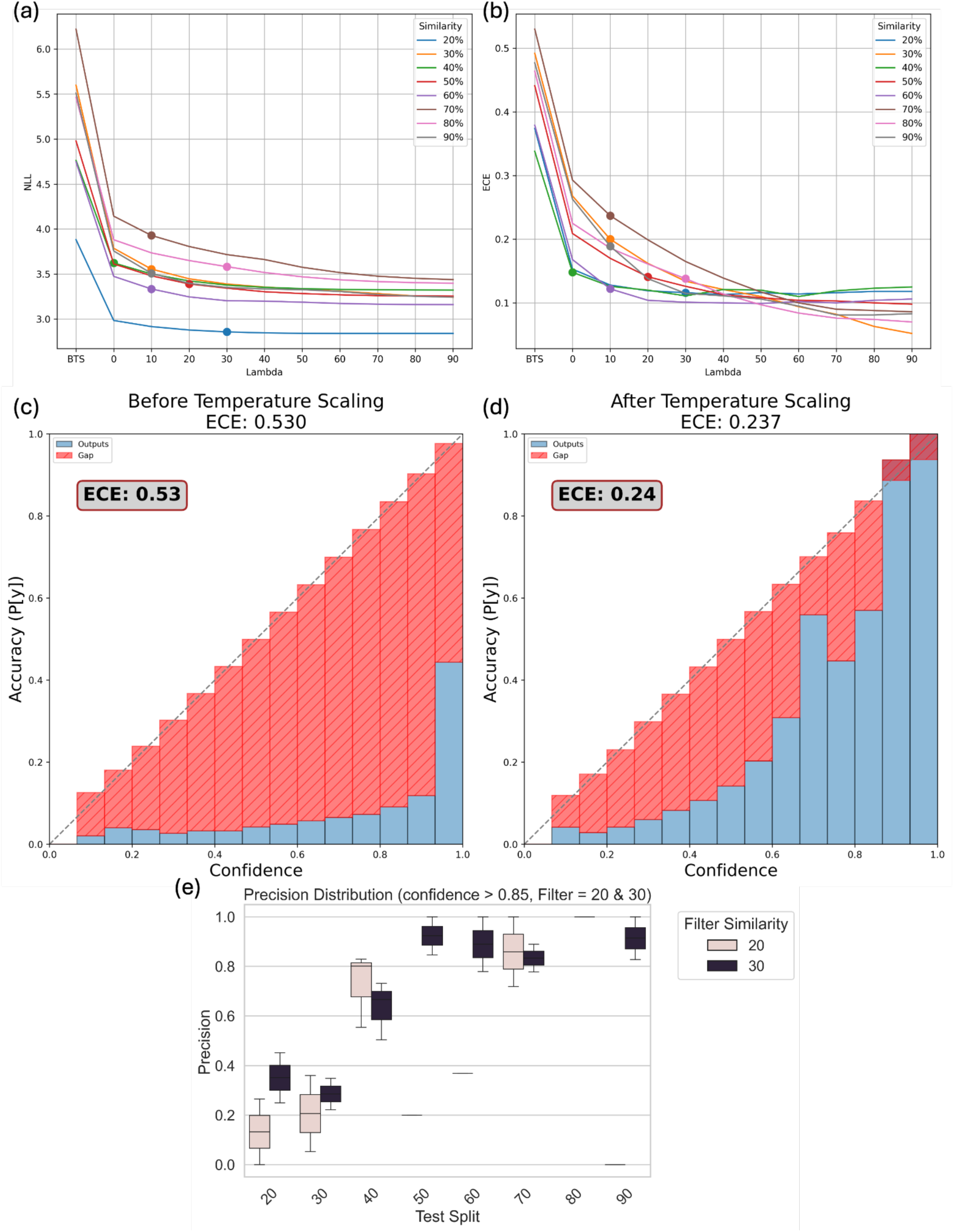
Diagnostic performance plots for BGF models undergoing temperature scaling. Temperature values were determined by minimizing a combination of negative log-likelihood (NLL) and a custom asymmetric loss function weighted by a scalar, λ (*see methods: temperature scaling*). Negative log-likelihood values (**a**) and expected calibration error (ECE; **b**) are graphically represented before temperature scaling (BTS) and across scalar values. For the λ value corresponding to the chosen final temperature for each model split, there is a point on each line. A reliability diagram shown prior to temperature scaling (**c**) and after temperature scaling (**d**) for the 70% trained model at λ=10 demonstrates calibration to a more realistic probability distribution. (**e**) Boxplot demonstrating the performance of model splits on the Filtered-Test set, at the 20 (brown box) and 30% (black box) percent identity threshold. Sequences were binned by their scaled softmax confidence output, and only those above a confidence of 0.85 were retained in bins of 0.05. The precision was computed for each of the bins and are represented by the boxes above. Some model splits only reported one populated bin above the 0.85 threshold, as represented by the black lines in place of boxes. Other model splits did not have any sequences above the softmax threshold for these splits (e.g., 20% filter identity for the 80% model), with no box present where this is the case.

Under any temperature scaling conditions, these metrics improved dramatically for all identity splits, with a continued improvement increasing the weight of the asymmetric loss function with increased penalty on overconfidence above 0.7. Calibration did not improve with higher identity splits (Fig 5a, 5b) as the 40% and 50% splits returned similar NLL and ECE scores as 90%. We also found that the best temperature did not necessarily correspond to the lowest ECE or NLL value, and deciding on a final temperature for each split was often guided by reliability diagrams (Figs S11-S18).

Using the 80% identity split as an example, expected calibration error continues to go down as the scalar λ increases (Fig 5b), however the softmax outputs become increasingly lower to the point of significant underconfidence in predictions with no predictions above 0.65 by λ = 50 or higher (Fig S17G). Overscaling resulted in a confidence cap where, while ECE and NLL were low, the model would not predict higher than a certain (often low) level of confidence (supplement). Through consideration of metrics and reliability diagrams for all identity splits prior we chose optimal temperature for each model. The resultant models can then be applied to unknown datasets while constraining the acceptable confidence associated with protein identification.

#### 2.2.2. After temperature scaling, certain model splits were able to confidently predict classes at distant identities using a high-confidence softmax threshold

As the overall goal of BGF is to identify unknown genes, we used the temperature-scaled model to identify a model split that provided the highest precision for remote homology. Through comparing the models that were trained using our range of identity scores and then applying them to both 20 and 30% identity splits, we identified that the model trained using the 70% test split with a softmax output above 0.85 resulted in precision between .75 and 1 which balanced loss of information with the risk of falsely identifying a significant portion of the genes (Figs 5e, S24). Or stated another way, when confronted with a protein that has less than 30% identity to the training set, the 70% BGF model alongside a confidence filter of 0.85 was able to identify genes with at least a .75 precision after the temperature scaling approach. The model trained at the 40% split performed relatively well compared to others as well, with a precision >0.5 for both 20 and 30% test splits (Figs 5e, S21). All other models were observed to have low precision for either one or both of the distant test splits (Figs 5e, S19-S26).

#### 2.2.3. Agreements across methods highlight the importance of a multi-method approach when exploring a cold seep metagenomic dataset

We then applied our temperature-scaled BGF model to a dataset of over six million protein sequences falling into over 3,000 unique MAGs from a global distribution of cold seeps. We found that there was significant intersection across all models, as well as certain model subsets (Fig 6). Across all of the methods a total of 1.05M sequences were annotated with BGF annotating 719,686, and HMMs the second most at 362,133. Additionally, these two approaches had the greatest number of predicted proteins that overlapped in their agreed-upon functions (Fig 6A, intersection size). BGF alone characterized 46% (485,849 sequences) of these proteins as belonging to the biogeochemical cycles that BGF was trained on. When contrasted with DIAMOND alignment to BGFdb or to KEGG, an often employed approach, 65,959 of the classifications were agreed upon across all of the methods. HMMs had the greatest number of annotations that were not in agreement with any other model (indicated by the solo point in Fig 6a), with BGF and BGFdb-DIAMOND following as the next-greatest in disagreement from other models.

**Fig 6.**
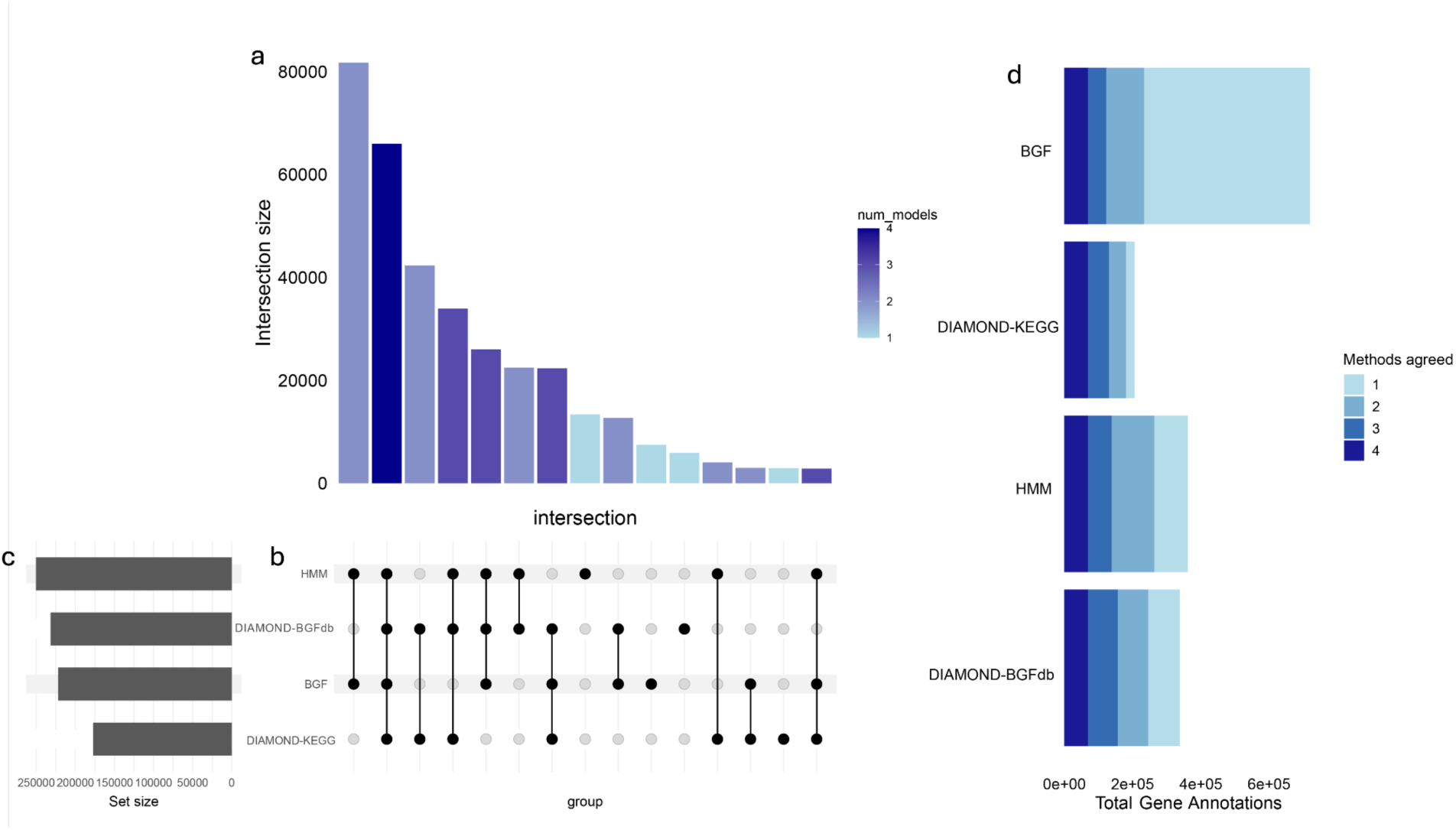
Upset plot showing the annotation agreement between models on a deep sea dataset comprising over 3,000 MAGs (*see methods: model application*). Intersection size (**a**) denotes the number of sequences agreed upon by a model-group combination (**b**). Set size (**c**) denotes the number of sequences for each model that fell into the majority of all models. Total gene annotations (**d**) represents the number of accepted gene annotations by each model on the dataset.

#### 2.2.4. BGF predicts more proteins related to nitrogen, methane, phosphorus, and sulphur cycling in certain pathways, with some pathways more affected than others

Comparing four different methods exhibits similar trends across the majority of biogeochemical pathways, however BGF classified substantially higher numbers of proteins for certain pathways (Fig 7). For example, when examining the serine cycling pathway, the alignment to BGFdb resulted in the greatest number of functional annotations (38,248 sequences), compared to other methods (BGF:16,902; HMMs:16,299; KEGG:20,641). In contrast, BGF predicted more proteins functionally related to organic phosphoester hydrolysis by orders of magnitude (43,564 sequences) compared to other methods (BGFdb: 1,342; HMMs:2,452; KEGG: 1,672). Another example with less extreme differences includes the central methanogenic pathway with more proteins predicted by BGF (33,250) compared to other methods (BGFdb:14,524; HMMs:11,540; KEGG:15,546), or denitrification (BGF:27,721; BGFdb:10,858; HMMs:710; KEGG:881). The two-component system and transporters pathways had significantly higher predictions by BGF and HMMs compared to alignment-based methods (BGF:113,751; BGFdb:13,206; HMMs:73,642; KEGG:4,100) and (BGF:38,708; BGFdb:11,841; HMMs:40,575; KEGG:6,589) respectively (Fig 7). Overall, Certain biogeochemical pathways including those related to phosphorus (two-component system, transporters, organic phosphoester hydrolysis, pyrimidine metabolism, pyruvate metabolism), methane (central methanogenic pathway, AOM) nitrogen (denitrification, assimilatory nitrate reduction, nitrogen fixation) appear to have greater potential of cryptic proteins participating in these pathways (Fig 7). Given that the HMMs constructed on BGFdb are overall high-performing at the 50% identity split (Fig 2), these model predictions can bolster the confidence in informing cryptic protein hypotheses complementing BGF’s predictions (two-component system, transporters; Fig 7).

**Fig 7.**
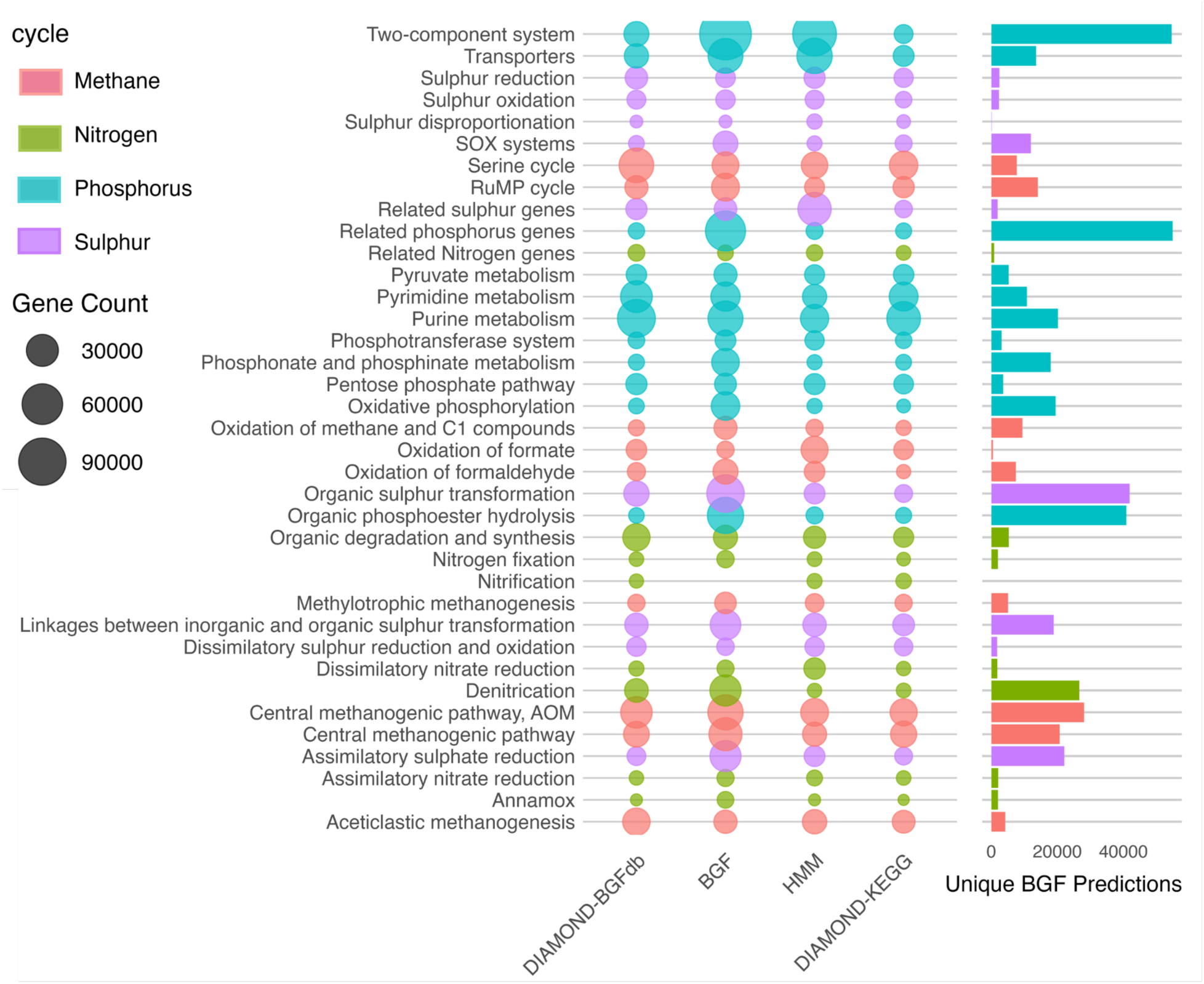
Bubble plot showing the magnitude of annotations for each prediction method into each class for the deep-sea MAG dataset. Protein count is represented by the size of each bubble, and bubbles are colored based on whether the metabolic pathway falls into methane cycling (red), nitrogen cycling (green), phosphorous cycling (blue), or sulphur cycling (purple). The bar plot on the right represents the number of predictions unique to BGF per pathway.

## 3. Discussion

### 3.1. Model refinement and advancement

BGF successfully classified proteins into 37 biogeochemical pathways for both similar and dissimilar proteins. A main component of BGF was shifting to define classes by function, and not sequence identity, which yielded a high-performing model and, critically, novel annotations (Figs 5,6,7). HMMs, in many cases considered the emergent leader in protein identification, were trained on this approach and BGF and HMMs trained at the metabolic pathway level identified a diversity of annotations shared with one another (Fig 6). While genes identified through both of these methods have only putative function until laboratory confirmation, overlapping predictions by diverse approaches (HMMs and BGF) provides some confidence in the identity of these heretofore uncharacterized proteins. BGF has a clear advantage over HMMs in its diversity of predictions as well as its performance metrics, likely due to the fact that classes were very broad and the MSAs constructed for HMMs were highly heterogeneous. However, BGF has less transparency than HMMs in how the predictions were made; transformers are often faulted as ‘black boxes’ which reduces the potential of discovering novel functional motifs that HMMs can extract [44]. Regardless, protein language models (PLMs) have become increasingly common and successful in biological applications [45–47], analogous to what we found through the training and application of BGF.

By applying a novel asymmetric loss function while temperature scaling, we were able to improve BGFs overall confidence to minimize erroneous protein function predictions (Fig 5). Given the importance of a trusted confidence threshold in bioinformatics, softmax values are used to represent true confidence. To calibrate softmax output, we applied and adapted [48]’s temperature scaling approach, adding an asymmetric loss function which penalized overconfidence. This novel approach is well-suited for balancing confidence calibration while also minimizing compute time required for model inference, enabling for the functional annotation of large ‘omics datasets within acceptable timeframes in contrast to other methods, including ensemble modeling, dirichlet scaling, and platt scaling [48–51]. These methods are more computationally costly, leading to longer run times in class prediction and show similar results in their ability to calibrate model output compared to a temperature-scaled softmax function [48]. Our asymmetric loss function improved ECE and NLL beyond classical temperature scaling, particularly when penalties are added above a 0.7 softmax threshold for incorrect predictions during model calibration (Fig 5).

Fine-tuning an ESM-2 model provides a forefront approach to leverage the power of a large-scale, generalized model on a narrow task [19,25]. ESM-2 models perform differently based on their size, however larger models have additional computational costs leading to a barrier in both training and wide-spread application. We chose the 8 million parameter model to optimize its efficiency as our goal was identifying protein function in large metagenomic datasets, and we found that it was high-performing, comparable to alignment approaches and better than HMMs while resolving significantly more genes. Scalability was a main criteria for our selection of models. As larger models can perform better on complex tasks, application of a larger model following the same training and testing set, and in particular our asymmetric temperature scaling, may provide additional insight into the function of earth system relevant proteins. A notable limitation is that ESM-2 models, including BGF, are limited by a context window of 1024 amino acids or less [19], complicating their ability to characterize larger proteins. As the field progresses using novel approaches, we can evaluate whether this limitation impacts the ability of the model to identify putative genes at remote distances past the abilities of alignment and HMMs, yet still functionally relevant.

### 3.2. Example Model Application

We chose cold seep ecosystems as a model to apply BGF as they have high microbial and functional diversity including a diverse suite of biogeochemical cycles; they also are a region of significant microbial dark matter in the oceans. Cold seeps are fueled by methane, providing the opportunity to better understand the genes and proteins involved in C_1_ cycling that are associated with greenhouse gas emission and mitigation, including taxa and genes currently unknown and likely involved [52–54]. Novel culture-free approaches such as BGF are therefore needed to address this research need in a multifaceted way using past and future ‘omics data. Further, these habitats have a suite of N-cycling, including nitrogen fixation and denitrification [55,56] that complement the well established role of S in the systems [57]. We show that cold seeps represent an excellent system in which deriving hypothetical gene function serves as a means for narrowing future research directions with pre-existing data. By annotating a large cold seep database composed of MAGs, we have shown the potential of BGF in identifying putative genes involved in a diversity of cycles, including in an environment often considered extreme. As an example of hypothesis generation, the number of additional genes predicted in the denitrification and assimilatory nitrate reduction pathways at cold seeps provide an impitus to apply experimental techniques to test the importance and coupling of N and CH_4_ oxidation, or highlight potential syntrophic relationships among taxa that carry out these functions. A surprising result was the prediction of significant numbers of cryptic phosphate cycling proteins, suggesting that direct study of these processes could be a fruitful direction of cold seep science. Other recent studies support this suggestion as well [26,58,59]. For example, LucaPCycle is a phosphorus-based PLM constructed and deployed on cold seeps, discovering 4,606 new phosphorus-cycling protein families [26]. A study of methane production in the water column provided evidence showing the importance of phosphorus in mediating methanogenesis [58], while other studies highlight the relative gap in knowledge around phosphorus cycling within cold seeps [59]. Overall, we show that BGF assists in hypothesis building and enables deeper exploration of ‘omics data generated from any ecosystem, linking proteins to biogeochemical cycling.

Much of the exploration of the vast, and ever expanding genomic data sets of critical ecosystems on earth is hampered by the unknown. No one approach will provide true insight into the genes, proteins, taxa, and processes that shape function, while at the same time, it is highly unlikely that the miniscule proportion of these genes that we ‘know’ the function of are entirely responsible for important processes. Our examination of the overlap of multiple models enhanced understanding and confidence in novel annotations (Fig 6). While no one approach is able to give a ‘true’ answer of gene functions, when used in ensemble they can provide a suite of novel insights into the genes, proteins and functions that together facilitate a diverse ecosystem functions, and especially those that current alignment-focused approaches miss [8]. BGF was able to highlight many genes that would otherwise be discarded, and in combination of alignment and HMMs, provide a wealth of insight and direction for future research on the realized function of earth systems. Through application of BGF, in concert with other approaches, to a diversity of earth systems we can make significant contributions to the landscape of genes, their diversity, and what drives ecosystem processes across the planet.

### 3.3. Conclusion

By fine-tuning a protein language model (BioGeoFormer or BGF), we explored microbial dark matter in proteins related to methane, nitrogen, sulfur, and phosphorus cycling. By using sequence identity clustering, we demonstrated the model’s ability to functionally annotate distant homologs, and compared this to alignment and HMM approaches. We highlighted the importance of uncertainty estimation and calibration, and applied temperature scaling to scale the softmax output to more accurately represent uncertainty, showing that the calibrated 70% identity model split outperforms all others in precisely annotating distant, yet functionally related proteins. Comparing our model to three other methods, we showed where BGF agrees with others, but also predicted novel putative function of currently unknown genes within a global cold seep dataset, highlighting further research directions in numerous biogeochemical pathways. Overall, we found that protein language models are an important tool in illuminating microbial dark matter, reducing unknown function, and supporting and directing future hypotheses in biogeochemical cycles as they relate to microbial function.

Given that some model splits outperformed the gold standard in remote protein prediction we recommend the use of the 70% identity split model, with a softmax confidence cutoff of at least 0.85 for sequences with less than 30% identity to the database considered. Therefore, we would like to emphasize that this new approach comes as a complement, not a replacement for the current gold-standard methods (i.e. > 30% identify, DIAMOND should be used). This method is for broad and hypothetical function assignment to drive future research directions, and should be used on sequences where high-performing, high-resolution methods such as alignment and HMMs do not return results.

## 4. Methods

### 4.1. Data curation, model training, and calibration

#### 4.1.1. Dataset curation

We combined four previously published datasets that grouped genes into distinct metabolic pathways: MCycDB (methane focused database [20,39–41]), SCycDB (Sulphur focused database [20,39–41]), NCycDB (Nitrogen focused database, [20,39–41]), and PCycDB (Phosphorous focused database [20,39–41]) with comprehensive methods for each database reported in their respective papers. Briefly, the authors of each database manually retrieved genes based on metabolic pathways in KEGG and public literature, and their respective protein families were downloaded from the UniProt database. Using the tool USEARCH [60] at a 30% identity cutoff, proteins from the UniProt sub-database TrEMBL were searched against the UniProt sub-database Swiss-Prot to form a “seed database.” This seed database was then searched, again using USEARCH at a 30% identity cutoff, against arCOG, COG, eggNOG, and KEGG, and homologous sequences were added to the database. Finally, corresponding sequences were added from the NCBI RefSeq database, which includes sequences from Archaea and Bacteria. To remove duplicate sequences from the fully constructed database CD-HIT was used at a 100% cutoff to ensure sequences were non redundant. We note that while KEGG served as the foundation for metabolic pathways within the CycDB databases, that KEGG pathways and CycDB pathways are not synonymous with one another. For example, the two-component system within KEGG includes genes associated with phosphate limitation, but also nitrogen regulation (ntrC family) and chemotaxis (cheA family). The two-component system in PCycDB, however, only includes the genes affiliated with phosphorus cycling [41]. To avoid any disagreement in contextualization and/or label application between the various databases employed or combined, we followed and adapted the CycDB databases, and its methods, when defining classes within BGF.

The goal of this study was to make broad putative functional annotations based on biogeochemical pathways and to assign protein families to their corresponding biogeochemical cycles. As a result, all protein sequences related to a particular cycle were grouped into a single class within BGF. For example, dissimilatory sulfate reduction within SCycDB [40] was considered a single class in training. An additional challenge is that certain proteins are known to be associated with more than a single biogeochemical function; to avoid leakage across classes, we filtered out sequences that fell into more than one class. A particularly challenging protein family is the central methanogenic pathway, where multiple proteins are associated with multiple processes, including the anaerobic oxidation of methane. Therefore, for biogeochemical processes associated with these two pathways, we classed unique proteins (“central methanogenic pathway”), as well as overlapping proteins (“central methanogenic pathway & anaerobic oxidation of methane (AOM)”). Ultimately, we had sequences grouped into 37 classes used for model training, validation, testing (Table S1). We heretofore refer to this curated and processed database as BioGeoFormer-db (BGFdb).

Using CD-HIT ([61]; version 4.8.1), each class within BGFdb was clustered at distinct sequence identities for 20% and 30% (psi-CD-HIT) as well as 40% to 90% at an interval of 10% (CD-HIT), and then split into a 60/20/20 scheme for training, validation, and test data respectively. Sequences were assigned to each dataset based on their cluster assignment. To quantify the relative performance of CD-HIT, we used DIAMOND ([62]; V 2.1.9.163) with -- best_hit to align test and validation datasets against the training dataset, reporting the median sequence identity between each class. These results were visualized and can be found within the supplementary information (Figs S1,S2).

We further processed the test set to evaluate each model’s ability in detecting distant, yet functionally related proteins. The greedy clustering algorithm in CD-HIT facilitated our goal of creating identity splits on large classes, yet led to “leaky” datasets where highly related sequences were maintained even after data splitting occurred (Figs S1-S4). Data leakage when using CD-HIT and similar tools is well documented [63,64], however more stringent methods such as hierarchical clustering used in previous studies leveraging PLMs proved infeasible due to the number of sequences within the highly populated classes (Table S1). Therefore, we used CD-HIT to perform clustering in order to understand model performance across a general trend of disidentity (Figs S1-S4). For more stringent analysis of each model’s performance at remote homologies, we combined the test set with the sequences dropped during clustering, and ran a DIAMOND alignment between training and test set. We then filtered the appended test set by the percent identity output from DIAMOND based on the model we were examining (e.g., the 30% model was evaluated with the 30% test set, which included sequences with 30% or lower percent identity between the train-test DIAMOND alignment). For sequences that had no match between train and test sets, we considered these to be below 20% identity and were labeled ‘unassigned’.

#### 4.1.2. Transformer model training

We used a pre-trained protein language model, ESM-2, that we tuned towards protein function to the pathways specified in BGFdb. [18,65]. While a family of ESM-2 models that range from 8 million to 15 billion parameters have been released, we used the smallest model, the 8 million parameter transformer with 6 layers, named ESM2-8m. In order to make this model perform sequence classification, we added a classification layer of 37 units (cycles). This layer varies between 35 and 37 units depending on the identity split. This allowed us to exploit the evolutionary representations that ESM-2 has learned from millions of sequences. All model training was done with HuggingFace [66]. We trained the model for 1 epoch with an initial learning rate set to 0.0003 with a linear learning rate schedule. We used the AdamW optimizer to train the model and a dropout probability of 0.1 across all fully connected layers in the model [67,68]. We used a batch size of 8 and trained the parameters in the ESM-2 backbone along with the added classification layer. We found that allowing all parameters to be fine-tuned resulted in the best performance. We trained a separate model for each identity split and utilized median sampling of the training set for a uniform distribution of cycles during training. The version of ESM-2 we used consisted of 6 encoder layers with a hidden dimension size of 1280 and output dimension of 320. The outputs from ESM-2 were pooled by using the CLS token position in the input sequence, which was added to the beginning of the input sequence. The embedding produced for this token was used as input to the classification layer to predict the biogeochemical cycle the input sequence belongs to. When training transformer models we also had to consider positional encodings for the tokens. In the case of ESM-2, the authors use rotary embeddings [69].

#### 4.1.3. Transformer model calibration

To ensure that the softmax output probabilities were as representative of true confidence as possible, we performed temperature scaling (equation 1). The inputs to the softmax function, or logits (z) were divided by a learned scalar parameter (T).

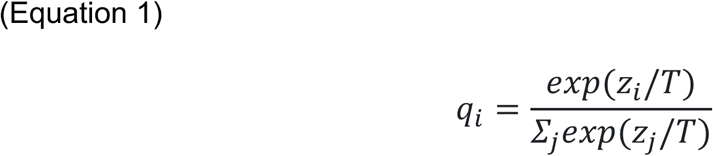

The scalar parameter (T) was determined by minimizing the loss on the validation set. To perform this, methods to minimize negative log likelihood (NLL) developed by [48] were used alongside a custom asymmetric loss function to minimize the overconfidence of the model when predicting false positives. Given that false positives are more costly than false negatives in biological research [70,71], we designed the asymmetric loss function to penalize overconfident, false positive predictions (equation 2), where *L*_*asym*_ is defined as the mean score across predictions, in which the value is above 0 only if the prediction is incorrect 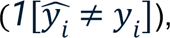 and only if the softmax confidence output is higher than a set threshold *t* (*max*(*0*, *c*_*i*_ − *t*)). When both of the above requirements are met, the difference between the confidence output and the threshold *t* are multiplied by a weight *w*. Therefore, the greater the confidence output is above the threshold, the greater the contribution to *L*_*asym*_.

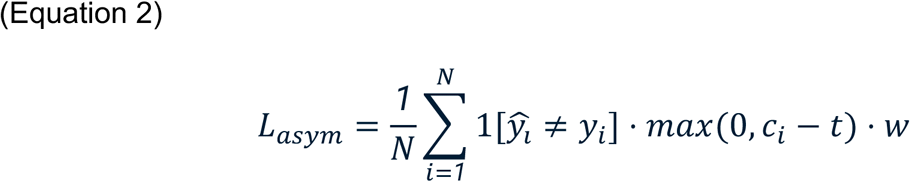

Temperature was calibrated by the L-BFGS optimizer over a maximum of 50 iterations. Values for the threshold *t* and weight *w* were set for equation 2 prior to optimizing (*t* = 0.7; *w* = 2.0). The temperature value was set by minimizing the negative log likelihood and the previously discussed asymmetric loss function (equation 3). The weight of the asymmetric loss function was manipulated by a scalar (*λ*) which was tested from 0 to 90 at intervals of 10 (equation 3). For each temperature value, the final negative log likelihood and expected calibration error were calculated and reported (Fig 5). To determine the final temperature for each model, NLL, ECE, and reliability diagrams were used in the decision making process.

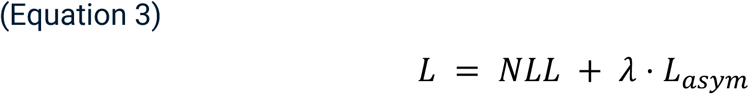

#### 4.1.4. Hidden Markov Model Training

To test the ability of commonly used remote homology detection methods on each class within BGFdb, we constructed hidden markov models (HMMs). Multiple sequence alignments (MSAs) were constructed using FAMSA (v. 2.2.2) for each class across all sequence identity splits for training data within BGFdb. HMMs were trained on each MSA using HMMER (v. 3.4) using default settings [44]. To evaluate the performance of HMMs, models were grouped by class and sequence identity, and evaluated on their respective validation and test sets. To directly compare against BGF, all HMM outputs trained at a specific sequence identity were treated as a cohesive “model” when computing performance metrics. The highest performing model was accepted as the correct classification based on the e-value followed by the bit score if there was an equivalent tie by e-values. No minimum e-value or bit score cutoff was used, in order to appropriately benchmark HMMs against BGF with no confidence cutoff.

#### 4.1.5. BGFdb alignment

To compare the performance of alignment methods on the training data, we used DIAMOND ([62]; V 2.1.9.163) to build a database for each identity split. We then ran DIAMOND with -- best_hit against respective validation, test, and filtered-test sets. Any match under these parameters was considered (no e-value cutoff) to appropriately benchmark against BGF with no confidence cutoffs.

#### 4.1.6. Performance metrics

For measuring performance of BGF, HMMs and alignment of BGFdb, we chose a set of metrics suitable for imbalanced multiclass classification. We used Matthew’s Correlation Coefficient (MCC) due to its relatively lower bias with varying class sizes [43,72]. We contrasted this performance metric with accuracy, precision, recall, and F1 score.

To observe what our model learned we extracted embeddings of all the test sequences from the hidden layer prior to the classification layer. We observed the community structure of proteins when we projected these embeddings into two dimensions using t-SNE [73]. Each sequence in the plot is colored by the biogeochemical cycles they were identified as belonging to. For each plot, 30,000 sequences were randomly selected from their respective test sets, perplexity was set to 30, and the learning rate was set to ‘auto’ with 1000 iterations.

### 4.2. Model application

#### 4.2.1. Dataset

We applied BGF to the metagenome-assembled genome (MAG) dataset constructed by [42], using 165 metagenomic samples from 16 cold seep sites around the globe. From this dataset, we used the 3,164 species-level MAGs constructed that included 81 ANaerobic MEthanotroph (ANME) species and 23 sulfate-reducing bacteria (SRB), among others. We predicted 6,282,958 proteins using Prodigal V2.6.3 [74], with an average 282.6 amino acids per protein (minimum 20, and maximum 17,512 amino acids [74,75]).

#### 4.2.2. Annotations

To functionally map proteins within the above dataset to our target biogeochemical pathways, we used and compared four complementary methods, ranging from sequence alignment to BGF’s deep learning approach. We used DIAMOND (v 2.9.4.115) ([62] to create a database from BGFdb (i), as well as to construct and run alignment on the KEGG “prokaryotes” database (ii). Both DIAMOND runs filtered annotations to return “best-hits” (e-value <= 1×10^-5^, percent identity >= 50, bit score >= 50). We additionally used the HMMs constructed at the 50% identity split of BGFdb following the HMM methods as used for model training (iii), returning “best-hits” (e-value <= 1×10^-5^, bit score >=25). We chose the 50% split because it performed above other HMM models (Fig 2). Last, we used BGF to predict protein function, using our model trained at the 70% identity split (iv) (*see methods: transformer model training*). We chose the 70% split because it was the most capably calibrated at distant identities, giving significantly more weight to its confident predictions for both similar and dissimilar proteins (Fig 5e, S24). We used a softmax probability cutoff of >0.85 to return high-probability classified proteins. Since deep learning allowed for pathway-level classification and alignment methods operate at the gene or KO level, we mapped alignment outputs to broader functional categories constructed based on genes within BGFdb to allow for a consistent and meaningful comparison.

To compare and contrast the annotations made by these four different methods on the same dataset we visualized their agreement and disagreement. To show the agreement between models on the cold-seep dataset, we used the UpSetR package [76]. From the cold seep dataset, proteins that had two or more annotations were retained for this visualization (Fig 6). To visualize the differences between models in the number of proteins predicted, all annotations were filtered to be within acceptable ranges of confidence per respective method. These were retained and the total number of annotations per method were visualized, alongside total annotations per class (Fig 7).

## Acknowledgements

We would like to extend our thanks to the Center for Quantitative Life Sciences (CQLS) as well as the College of Engineering at Oregon State University for providing the computing resources necessary to complete this project. We would also like to thank Dr. Christine Tataru for participating in early discussions and ideation while conceptualizing the project.

## Funding

Funding for this work was provided by the U.S. National Science Foundation RISE - 2425835, OPP-2046800 to AR Thurber, NSF - GRFP 2139319 to JH Wynne, CAIG - 2425834 to MM David.

## Data availability statement

There are no primary data in the paper; all scripts used to reproduce this study are available on Github (https://github.com/nimuh/biogeoformer/tree/main). Due to space limitations, the GitHub repository does not have many files used to conduct the study. The completed repository is available for download at Zenodo [77].

## Notes

### Competing Interest Statement

The authors have declared no competing interest.

